# GelMA as scaffold material for epithelial cells to emulate the small intestinal microenvironment

**DOI:** 10.1101/2024.06.24.600349

**Authors:** Inez Roegiers, Tom Gheysens, Kim Vanbeversluys, Nikoletta Rać, Grzegorz Stroka, Jana de Croock, Tom Van de Wiele, Peter Dubruel, Marta Calatayud Arroyo

## Abstract

Host-microbe interactions in the intestine play a significant role in health and disease. Novel scaffolds for host cells, capable of potentially supporting ese intricate interactions, are necessary to improve our current systems for mimicking host-microbiota interplay *in vitro/ex vivo*. In this research paper, we study the application of gelatin methacrylamide (GelMA) as scaffold material for intestinal epithelial cells in terms of permeability, mechanical strength, and biocompatibility. We investigated whether the degree of substitution (DS) of GelMA influences the permeability and found that both high and low DS GelMA show sufficient permeability of biorelevant transport markers. Additionally, we researched epithelial cell adherence and viability, as well as mechanical characteristics of different concentrations of GelMA. All concentrations of hydrogel show long-term biocompatibility for the mono- and co-cultures, despite the goblet-like cells (LS174T) showing lower performance than enterocyte-like cells (Caco-2). The mechanical strength of all hydrogel concentrations was in a physiologically relevant range to be used as scaffold material for intestinal cells. Based on these results, we conclude that GelMA is a suitable material as a scaffold for intestinal cell types in terms of permeability, mechanical strength and biocompatibility. These findings contribute to the growing field of *in vitro* modeling of the gut and moves the field further to ensuring more translatable research on host-microbe interactions.

## 1. Introduction

The intestinal epithelial surface is a key defensive barrier that protects us from the outside world, continuously interacting with commensal bacteria, potential pathogens and dietary components. The small intestinal microbiota, epithelium, the layers underneath the epithelium and the interactions occurring between previously mentioned components are involved in several physiological processes driving immune development, modulating metabolic homeostasis or promoting colonization resistance against pathogens ^1,2^. Sampling procedures to retrieve human native intestinal tissue or microbiome samples are invasive and expensive, and animal models typically offer only limited human translatability. This is limiting our understanding of the dynamics and behavior of microbial populations in their interaction with human cells ^3–5^.

In response to these challenges, three-dimensional (3D) human models have emerged as a new paradigm that can accurately replicate complex physiological responses *in vitro* ^6^. These advanced models are based on progress in cell biology, micro-engineering, biomaterials, and biofabrication^7^. Emphasis is placed on developing innovative and stable scaffold materials that mimic an extracellular matrix (ECM), which facilitates the adhesion and growth of – in this case – small intestinal epithelial cells. The ECM serves as a physical support and regulates cellular processes such as differentiation, adhesion, and migration through various biochemical cues. It comprises water, proteins, glycoproteins, and polysaccharides, forming a three-dimensional network. The ECM and cells display a two-way communication, and the matrix is constantly remodeled by cells, making it a dynamic system^8^.

For *in vitro* applications, hydrogels form an excellent material to mimic the ECM. Hydrogels possess the potential to mimic cell microenvironments due to their flexible properties. These macromolecular networks have a hydrophilic nature and can incorporate a high water content, which is vital for nutrient and residue transport, supporting cell survival (reviewed by^9^). Gelatin, a biomaterial obtained from the denaturation of collagen, is considered highly interesting for making hydrogels as it contains the arginine-glycine-aspartic acid sequence (RGD), which is the most common peptide motif responsible for cell adhesion to the ECM^10^. The utilization of collagen gels as a scaffold material for various cells has been thoroughly researched. Particularly in the context of the intestine, extensive research has been focused on investigating the integration of intestinal and primary stem cells on 3D collagen scaffolds, which were fabricated prior to cell seeding by molding or microfabrication techniques to accurately replicate the geometric characteristics of intestinal villi ^11,12^. Li and colleagues have utilized collagen gels to seed fibroblasts, Caco-2 cells, and HT29-MTX cells, creating a 3D triple co-culture model ^13^. This model has been successfully employed to assess drug permeability and has demonstrated drug absorption rates that are more representative of physiological conditions ^14^. Often, materials used as scaffold for intestinal epithelial cells such as polyethylene terephthalate (PET) and polycarbonate (PC) are coated with a commercially available ECM-like substance such as a collagen type I coating, MaxGel™ (human-derived) and Matrigel (mouse-derived)^15–18^. We would like to simplify this by proposing the use of a cost-effective, biocompatible and relatively easy-to-handle hydrogel that does not require coating, that has the potential to be 3D-printed to obtain the characteristic microarchitecture of the small intestinal microenvironment, such as what is being presented in this research paper.

Since Van Den Bulcke and colleagues introduced methacrylamide side groups to gelatin to produce gelatin methacrylamide (GelMA), they introduced the ability to crosslink gelatin and making it more stable for long-term applications and thus providing a broader application range^19^. GelMA is an attractive biomaterial due to its peptide backbone providing suitable cell adhesion sites, and its ability to undergo photopolymerization to form covalently crosslinked networks. This method of crosslinking gelatin is a UV-based system which allows for the fine tuning of the degree of crosslinking by tuning the UV light, the photoinitiator type and concentration and the acrylate content. GelMA has already been used as scaffold for several different cell types, which is indicative of its potential (Table 1 ^20–25^). This wide application together with the long-term stability at physiologically relevant temperatures, low production cost and wide availability, excellent cell compatibility and no need for coating with ECM-like materials, makes GelMA an excellent candidate for a scaffold for intestinal epithelial cells.

**Table 1.** Culture time of several cell types on GelMA scaffolds.

| Cell type | Culture time on GelMA (Days) | Reference |
| --- | --- | --- |
| Human foreskin derived fibroblasts (HFF) | 7 | 21 |
| Primary human bone marrow derived cells (MSCs) | 7 | 20 |
| Human umbilical vein endothelial cells (HUVECs) | 7 | 22 |
| Human dental pulp stem cells (hDPSCs) | 7 | 23 |
| Human adipose-derived stromal stem cells (ASCs) | 14 | 24 |
| Intestinal epithelial cells (Caco-2) | 14 | 25 |

The effects of GelMA on cell phenotype, endothelial differentiation, cell proliferation, and migration have been demonstrated for several cell types^23,26^. These effects can be finely adjusted through various parameters of crosslinking, such as initiator concentration, GelMA methacrylation degree, macromer concentration, UV exposure, and curing time ^7^. Recently, Pamplona et al. investigated GelMA hydrogels with tunable mechanical properties for culture of Caco-2, generating GelMA scaffolds with different stiffness and testing the suitability of the materials as scaffolds for intestinal cells and looking at cell viability until 14 days after seeding^25^. Vila and colleagues constructed a GelMA-poly(ethylene glycol) diacrylate (PEGDA) hydrogel co-network that provides the required mechanical and biochemical features to mimic both the epithelial and stromal compartments of the intestinal mucosa in which fibroblasts were embedded, and Caco-2 cells were seeded on top ^27^. Their intestinal mucosal model was able to mimic some of the features attributed to mesenchymal-epithelial interactions such as enhanced epithelial cell proliferation and barrier permeability, diffusivity properties that allow paracrine effects and accelerated tight junction recovery. Szabo et al. created a 3D-printed GelMA-AEMA scaffold with intricate villi structures, on which they seeded a Caco-2/HT29-MTX coculture. They found that the 2D gel-MA-AEMA scaffold enabled the formation of a functional cell monolayer after 21 days ^28^. Our results show that the GelMA scaffolds can sustain a stable co-culture with the two most common cell types of the intestine, enterocytes and mucus-producing goblet cells, thereby confirming earlier findings as the GelMA being a suitable scaffold material for cells.

In this research paper, we investigate the potential of GelMA as scaffold material for intestinal epithelial cells. Firstly, to guarantee that small molecules can pass through the GelMA without the scaffold being the limiting factor, so that in the future, it can be applied for the study of host-microbe interactions in for example a transwell set-up (e.g. epithelium and microbiota on apical side, immune cells on the basal side, or variations on this set-up), we tested the diffusion of a range of biorelevant transport markers through the membrane while varying the degree of substitution (DS). After this, we varied the hydrogel concentrations, of which we tested mechanical strength and long-term cell adherence and viability. This work adds to the current research on novel scaffold materials for intestinal models and provides more insight in the potential of the scaffold enabling host-microbe interactions.

## 2. Materials and methods

### 2.1. Synthesis GelMA

Gelatin (type B) isolated from bovine skin by the alkaline process was supplied by SKW Biosystems, Ghent, Belgium. Methacrylic anhydride (MAA) was obtained from Aldrich (Bornem, Belgium) and was used as received. Dialysis Membranes Spectra/Por 4 (MWCO 12000−14000 g mol^-1^) were obtained from Polylab (Antwerp, Belgium). 2,4,6-Trinitrobenzene-sulfonic acid (TNBS) analytical grade was purchased from Serva (Heidelberg, Duitsland) and sodium azide from Acros Organics (Pittsburgh, PA).

GelMA was obtained following a previously reported protocol by Van den Bulcke et al. (Van Den Bulcke et al., 2000b). Gelatin methacrylamide was prepared by reaction of gelatin with methacrylic anhydride. The degree of substitution (DS) is defined as the percentage of ε-amino groups that are modified. Gelatin methacrylamide with a range of degrees of substitution were prepared by analogous synthesis by changing the amount of methacrylic anhydride. For this research, GelMA with a low DS (66) and high DS (91.43-98) was made and used for the diffusion cell experiments. After dissolution of gelatin in 0.1M phosphate buffer (pH 7.8) at 40 °C, methacrylic anhydride (5.74 mL for the low DS (1 eq), 14.34 mL for the high DS (2.5 eq)) was added while vigorously stirring. After 1 h of reaction, the reaction mixture was diluted with 1 L double distilled water and dialyzed (MWCO 12000−14000 g mol^-1^) for 24 h against distilled water at 40 °C, with a total dialysis volume of 60L and without any flow. After dialysis, the pH of the solution was adjusted to 7.4 using 0.1M NaOH. The solution was then distributed over several petri dishes and allowed to gellate at RT, whereafter they were frozen at - 20 °C. This could then be freeze-dried at -80°C and 0.37 mbar, leading to a white solid (Christ freeze-dryer alpha I-5). The degree of substitution (DS) of the High DS GelMA was determined using ^1^H-NMR spectroscopy and OPA-analysis. To determine the modified versus non-modified lysines, a proton nuclear magnetic resonance (*^1^H-NMR*)-analysis was performed on the GelMA. Of the GelMA sample, 10 mg was measured in a small glass vial. This was dissolved in 1 mL of deuterated water (D_2_O) at 40°C on a shaker. When the GelMA was completely dissolved, 750 µL of the solution was transferred to an NMR tube. The sample was analyzed with a 500 MHz NMR spectrometer (Bruker) at 40°C. For the OPA-analysis, This protocol was based on what was previously described by Van Vlierberghe and colleagues^29^. O-phthalaldehyde spectrophotometric (OPA)-analysis was conducted to determine the reaction of OPA and 2-mercaptoethanol with amino groups, which measures proteolysis and thus gives an estimation of the DS of the GelMA.

### 2.2. Synthesis of Lithium (2,4,6-trimethylbenzoyl)Phenylphosphinate (Li-TPO-L) (Photoinitiator)

In short, photoinitiator lithium (2,4,6-trimethylbenzoyl)phenylphosphinate (Li-TPO-L) was prepared: 8.60 g (27.2 mmol) of (2,4,6-trimethylbenzoyl)-phenyl-phosphinic acid ethyl ester (commercial Speedcure TPO-L from Lambson) and 9.45 g (109 mmol) lithium bromide were dissolved in 150 mL butanone and stirred for 24 h at 65 °C. The resulting precipitate was collected by suction filtration, washed with petrol ether, and dried under vacuum at room temperature.

### 2.3. Hydrogel synthesis

The hydrogels were prepared by radical crosslinking of solubilized gelatin derivatives. Crosslinking of the methacrylamide modified gelatin was performed in aqueous medium in the presence of a water-soluble photoinitiator, Li-TPO-L. The mixture was then poured into a cast made of two UV-permeable plexiglass plates separated by a silicon spacer with 1 mm thickness. This was then put at 4°C for 30min for physical gelation, since according to the upper critical solution temperature (UCST) behavior of gelatin, cooling the samples before UV crosslinking promotes the formulation of a physically crosslinked polymer network and therefore has a strengthening effect on the samples. Afterwards, the samples were immediately cured with UV-A light for 30 min (365 nm, 10 mW/cm^2^) using the VL-400L model (Vilber Lourmat, Marne La Vallée, France). After removal of the plexiglass plates, a flexible 1 mm thick transparent hydrogel film was obtained. For the diffusion experiments, disks with a 1.4 cm diameter were punched and sterilized by putting the samples in EtOH (70 v/v%) for 24h and afterwards exposing the samples to UVC for 30 min. For the cell adhesion experiments, the disks were sent for cold-cycle ethylene oxide sterilization (Animal Research Centre KU Leuven, Leuven, Belgium). Before use for the diffusion and cell adhesion experiments, the disks were equilibrated for 24h in cell culture medium (DMEM) at 37°C.

### 2.4. Gel fraction and mass swelling ratio of the hydrogel disks

The methodology to determine the gel fraction and mass swelling ratio has been used extensively ^28,30^. To determine the gel fraction, the crosslinked hydrogel disks (diameter = 8 mm, thickness = 1mm) were freeze-dried determining the dry mass (m_d1_). Next, the dried hydrogel disks were incubated in ddH_2_O at 37 °C for 24 h, followed by freeze-drying to determine the second dry mass (m_d2_). The gel fraction was determined using equation (1).

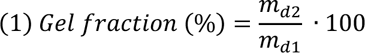

To determine the mass swelling ratio, the hydrogel disks (diameter = 8 mm) were incubated immediately after crosslinking for 24 h in ddH_2_O at 37 °C to obtain equilibrium swelling. Next, the hydrated mass of the samples was measured (m_h_) and the samples were lyophilized to determine their dry mass (m_d_). The mass swelling ratio was determined using equation (2).

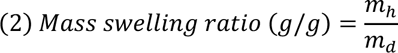

### 2.5. Rheological measurements to determine storage modulus (G’) and loss modulus (G’’)

Dynamic shear oscillation measurements at small strain were used to characterize the viscoelastic properties of cross-linked methacrylamide-modified gelatin hydrogel films. These rheological measurements at oscillatory shear deformation were carried out with a type Physica MCR301 (Anton Paar, Sint-Martens-Latem, Belgium) using parallel rough plates of 40 mm diameter and plate-to-plate distance of 900 µm. Mechanical spectra were recorded in a constant strain mode, with a low deformation of 0.05 maintained over the frequency range of 0.01−10 Hz (rad/s) at 37°C. The diameter of the plates was 40mm, compared to the hydrogel disks having a diameter of 12 mm.

### 2.6. Diffusion cell set-up and measurements of fluorescent compounds

Diffusion of three fluorescent molecules [(1) Lucifer yellow (LY) (Merck), (2) Fluorescein isothiocyanate–dextran average mol wt 4000 (FITC-Dextran 4, D4K) (Sigma-Aldrich) and Fluorescein isothiocyanate–dextran average mol wt 10,000 (FITC-Dextran 10, D10K) (Sigma-Aldrich)] was evaluated by placing a membrane in a side-by-side diffusion set up. To this end, a 250 µM solution in DMEM was prepared with either LY, D4K, or D10K. The setup is built up out of two diffusion cells, each diffusion cell is supplied with a stirring bar and the temperature is kept at a constant of 37 °C. The produced GelMA membranes (disks of 1mm thickness and 1.4 cm diameter, sterilized by 24h of 70% EtOH followed by 30min of UVC, equilibrated in DMEM for 24h) were clamped between the diffusion cells. The fluorescent compound was added at the apical side of the cell. 10% of the volume of the acceptor cell (300 µL) was emptied at each sampling point (30, 60, 90, 120, 180, 240, 360 and 1140 min) to measure fluorescence.

Fluorescence was measured in three technical replicates in black fluorotrac 96-well plates with flat-bottom (Greiner Bio-One) at an excitation wavelength of 540 nm and an emission wavelength of 590 nm for LY, and an excitation wavelength of 490 and emission wavelength of 520 nm for D4K and D10K.

### 2.7. Determination of Permeability Coefficient of Permeability Marker Compounds calculations

The apparent permeability coefficient (P_app_, cm/s) over 360min for each marker compound was calculated according to equation (3):

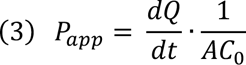

where dQ/dt is the rate of compound transfer (pmol/s) into the basolateral compartment under sink conditions (where less than 20% of the compound was transferred across the cell monolayer), A is the surface area of the filter insert (cm^2^), and C_0_ is the initial concentration in the donor chamber.

### 2.8. Cell culture

The reagents used in the following sections were purchased from MilliporeSigma (Burlington, MA, USA), unless otherwise stated.

In this study, two immortalized cell lines were used. Caco-2 cells were combined with LS174T cells, a mucus-secreting cell line, both derived from a human colorectal adenocarcinoma. The immortalized cell line Caco-2 (ECACC 86010202) was obtained from the European Collection of Authenticated Cell Cultures, (Public Health England, London, United Kingdom) and LS174T (CLS 300382) cell line was obtained from CLS (Cell Line Service, Eppelheim, Germany). When co-cultured, the Caco-2 cells and LS174T were combined in a 9:1 ratio resp. to mimic the *in vivo* distribution of cell types. T Caco-2 and LS174T cells (passage number, 7–35) were routinely grown as described with high glucose (4.5 g/L) and 1% (v:v) GlutaMax (Thermo Fisher Scientific), supplemented with 10% (v:v) heat-inactivated fetal bovine serum (iFBS; Greiner Bio-One, Kremsmunster, Austria), and 1% (v:v) penicillin/streptomycin/amphotericinB (ThermoFisherScientific). Caco-2 cell medium was also supplemented with 1% nonessential amino acids (Thermo Fisher Scientific). Cells were seeded with a density of 300.000 cells/mL DMEM, the total volume that was seeded in all conditions was 1.5mL. In brief, Caco-2 and LS174T cells were independently maintained following routine cell culture techniques. To conduct biocompatibility assays, cells were trypsinized, counted, and seeded in a co-culture of mixed cells (300.000 cells/mL; ratio Caco-2/LS174T = 90/10) (Fig. 1).

**Figure 1.**
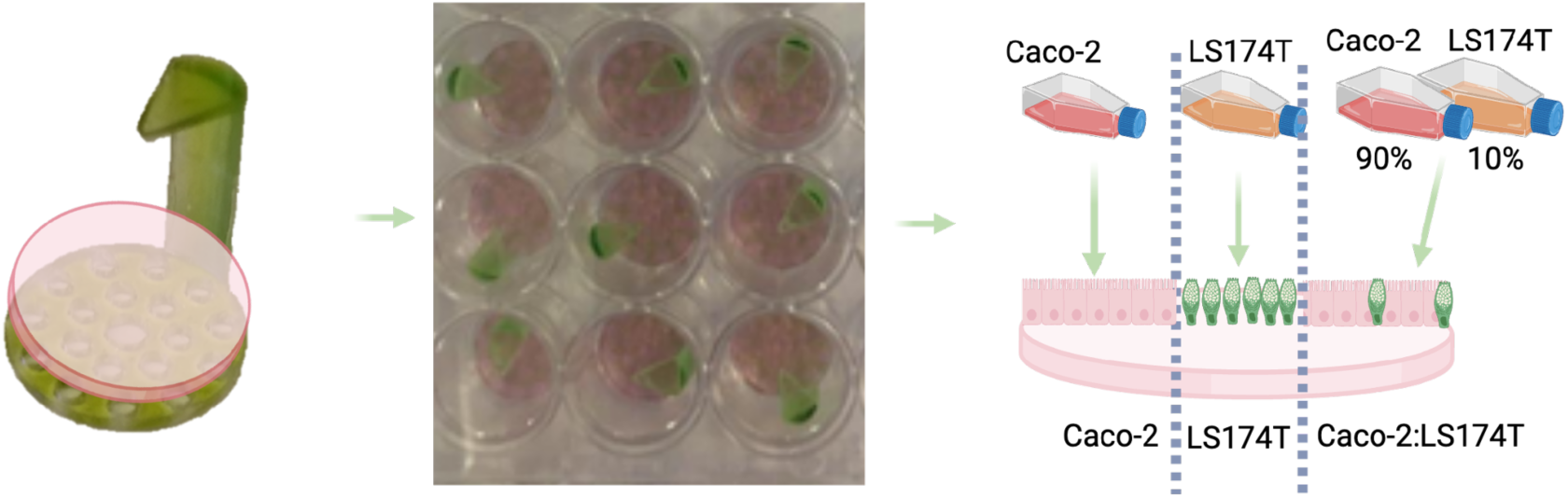
Schematic of seeding process of Caco-2, LS174T and the co-culture on the Gel-MA scaffold. Left: Crosslinked GelMA hydrogel on top of 3D-printed insert. Middle: Hydrogel disks on hydrogel insert were placed in 24-well plate. Right: Cell seeding of either Caco-2 or LS174T monoculture, or a co-culture of both cell types by pipetting cell suspension in DMEM on hydrogel disks on top of inserts.

The disks were put on 3D printed inserts made from Monomer Resin (Anycubic translucent green resin, Anycubic Photon SLA printer, Anycubic, Shenzhen, Guangdong, China) (Fig 1 and Fig S1 and S2). Post processing was done by washing 6 minutes in acetone after which the insert was dried at RT. Once dry, the insert was put on a rotating platform in a UV chamber with LED light (410 nm) (Anycubic Wash and Cure 2.0). Afterwards, the inserts were put into 2 liters of distilled water for 2 days for removing any residuals. The inserts were sterilized by submerging them for 24h in 70% EtOH followed by exposure to UV-C for 30 min. The insert material was tested for Caco-2 and LS174T adherence using the resazurin assay described in section 2.9. We observed that cells adhered to the insert material after initially seeding the cells on the hydrogel on top of the insert. After the cells adhered to the hydrogel, and any loose cells were washed away when refreshing medium 24h after seeding, and then transferring the hydrogels to a new, sterile 3D-printed insert, no cells adhered to the insert itself. To be completely sure that the inserts were not interfering in any case with the resazurin assay results, each time before metabolic activity was measured, the hydrogel disks with cells were therefore transferred before adding the resazurin to a new, sterile, insert to completely ensure only the metabolic activity of the cells on top of the hydrogel was measured. The control condition was cells seeded in the same density on the bottom of a 24-well plate treated for cell-culture (Corning Incorporated, New York, USA).

### 2.9. Metabolic activity

Living cells maintain a reducing environment within their cytoplasm and mitochondria, in which resazurin (blue and non-fluorescent) is reduced by dehydrogenase enzymes to form the red fluorescent dye resorufin. The amount of resorufin can be monitored by measuring fluorescence or absorbance, which is proportional to the number of living cells (“viability” or “metabolic activity”) in the sample. Metabolic and mitochondrial activity of host cells was assessed with a resazurin assay (7-Hydroxy-3H-phenoxazin-3-one-10-oxide sodium salt, Sigma-Aldrich, Overijse, Belgium). Medium was refreshed with 1 mL of 0.01 mg/mL resazurin in DMEM and incubated for 2 h at 37°C, 10% CO_2_ and 90% relative humidity. Fluorescence was measured in technical triplicates in black 96-well plates at an excitation wavelength of 540 nm and an emission wavelength of 590 (SpectraMax M2 plate reader, Molecular Devices, Brussels, Belgium). As a negative control, the fluorescence of resazurin in DMEM incubated in the absence of host cells was subtracted as a background from all fluorescent values. Metabolic activity was normalized to surface area where the cells were growing on (1.9 cm^2^ for the control, and 1.6 cm^2^ for the hydrogel disks). Metabolic activity results displayed in Fig. 3 and Fig. S6 for each cell type of the co-culture are results from one experiment, done in triplicate.

### 2.10. Staining and Confocal microscopy

For immunofluorescence staining, the cells were washed 2 times with PBS and fixed with 4% paraformaldehyde (Carl Roth, Karsruhe, Germany) for 10 min at room temperature. After 2 washes with PBS, cells were permeabilized with 0.1% Triton X-100 in PBS for 5 min and filamentous actin cytoskeleton staining was performed using rhodamine–phalloidin stain (MilliporeSigma) diluted 1:100 (v:v) in 0.5% normal goat serum (MilliporeSigma). The coverslips were mounted in Vectashield antifade mounting medium (Vector Laboratories, Burlingame, CA, USA) containing 1 mg/ml DAPI (MilliporeSigma) to evaluate the nuclear morphology. Visualization was performed on an Axioskop fluorescence microscope (Zeiss) and on a Nikon AR1 confocal microscope (Tokyo, Japan).

### 2.11. Statistics

All statistical analyses were performed using R version 4.0.2 (© 2020 The R Foundation for Statistical Computing). The package ggplot2 was used for plotting. Normality of the data was assessed using the Shapiro-Wilk test and homoscedasticity was assessed with Bartlett’s test for equal variances. In case normality and homoscedasticity were rejected, the Kruskal-Wallis test was used for multiple comparisons (K-W), followed by Dunn’s post hoc test with Holm’s correction, or for comparisons between two groups the Mann-Whitney U test was performed (M-W U). When normality and homoscedasticity were not rejected, two-way ANOVA was performed, followed by Tukey HSD post hoc test. Significance was considered at the 5 % level (α = 0.05). If the test is not specifically mentioned after a p-value, a parametric test was performed. Non-parametric tests are specifically mentiond as either K-W or M-W U). The significance level is also shown by following symbols for following results: *, **, *** and ****for p ≤ 0.05, p ≤ 0.01, p ≤ 0.001 and p ≤ 0.0001 respectively.

## 3. Results and discussion

### 3.1. GelMA high and low DS is equally permeable, both materials show adequate permeability to be used as scaffold material for small intestinal cells

The ability for small and large molecules (e.g. short-chain fatty acids, free fatty acids, amino acids and vitamins by the host) to go through a scaffold material is crucial when creating a mimicking environment for host-microbiota interactions studies ^31,32^. To ensure that the GelMA biomaterial does not hinder the transport of specific molecules across the epithelial barrier, we assessed the paracellular permeability of LY, and FITC labeled dextrans (4 and 10kDa) through the GelMA membranes with either a high or low degree of substitution (DS). The DS is important as it could directly impact the hydrogel pore size and thus, permeability ^33,34^. In permeability assays of epithelial cell layers, these compounds are commonly used and known to undergo paracellular transport through adherens and tight junctions located in between the epithelial cells, which is a diffusion-driven process across a concentration gradient over the epithelium. We calculated the apparent permeability (P_app_) values over 6h (the linear phase of diffusion) from apical to basal side (A to B), shown in Figure 2. For LY, D4K and D10K, the observed values are in the range of 10^-6^-10^-7^ cm/s. For LY, we observed a P_app_ of 2.88·10^-5^ ± 7.16·10^-6^ cm/s and 3.02·10^-5^ ± 4.47·10^-6^ cm/s for Low DS and High DS respectively (mean ± St.dev), with no significant differences between the two DS’ (M-W U). For D4K, observed values were 5.83e-06 ± 8.60e-07 and 7.79e-06 ± 5.29e-06 for low and high DS resp. wit no significant differences between the two DS’ (M-WU), and for D10K values were 4.74e-6 ± 2.53e-06 and 3.63e-06 ± 7.61e-07 with no significant differences between DS’ (M-W U).

**Figure 2.**
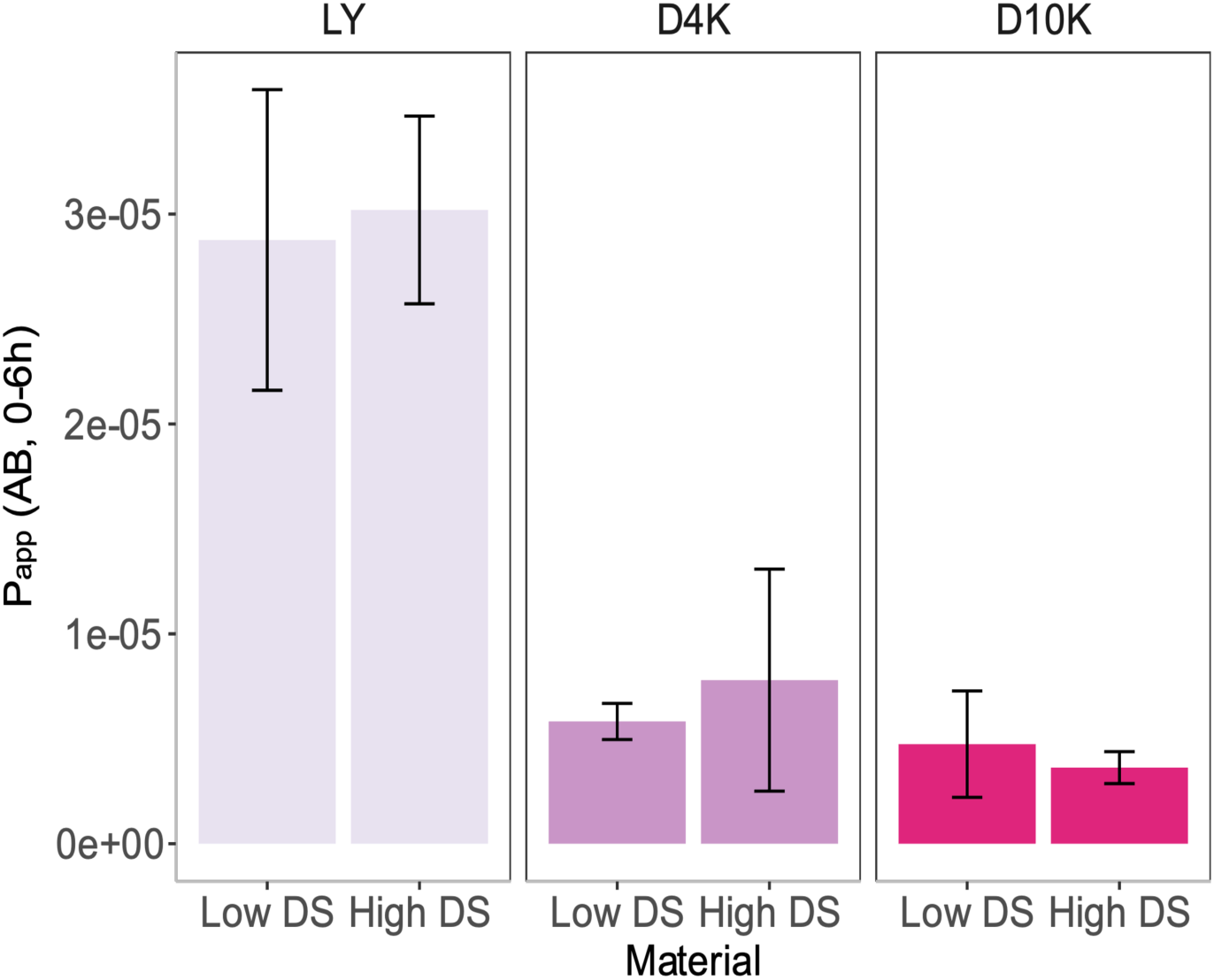
Apparent permeability (P_app_) (Apical to basal, abbreviated as AB) after 6h for Low DS and High DS GelMA, and the silk membranes. Tested compounds are Lucifer Yellow (LY); FITC-Dextran 4kDa (D4K) and FITC-Dextran 10kDa (D10K). Bars represent the average P_app_ ± standard deviation (n=2 except for D4Kwhere n=3). No significant differences in Papp over 6h were found in between materials for each compound (M-W U)

The results indicate that the selected compounds exhibit equal permeability across all tested materials. Notably, the permeability values obtained in this study for scaffolds intended for Caco-2 cell culture in the small intestine are approximately 10 times higher compared to values reported in literature (on scaffolds with a confluent layer of Caco-2 cells), suggesting that the scaffold material is suitable for this application^35–41^. It is worth noting that values in literature are reported for membranes with Caco-2 cells, which are cells that form a very tight monolayer. Here, the permeability was not tested with a cell monolayer due to practical limitations, yet this could be tested in the future, perhaps in a more advanced set-up suitable for cell cultures such as the Ussing chamber^42^. It should also be noted that the P_app_ calculated from transwell experiments cannot be directly compared to the P_app_ calculated from this experiment, due to it being a different set-up (the diffusion set-up used in this experiment has constant stirring, whereas transwell experiments are mostly static). Previous studies have reported permeability of specific compounds in Caco-2 stirred monolayers to avoid the effect of unstirred water layers in permeability tests, however no direct comparison with LY or dextrans can be performed^43,44^.

In the context of using GelMA as a drug releasing vehicle, Shao and colleagues found that GelMA can interact with drug by physiosorption and covalent linking. In general, drug delivery from GelMA is mediated by diffusion and degradation. They found that the molecular weight of drugs and the pore size of GelMA play important roles in the release process^45^. Kim and colleagues showed that gel integrity and gel drug binding can have profound effects on the release of drugs from hydrogels over time^46^. This is relevant if the GelMA would be used as scaffold material for intestinal epithelial cells in an *in vitro* model to research, for example, drug interactions with the host, and would require more research in how other compounds interact with GelMA. It is likely that other factors like the degree of crosslinking of the hydrogel, the presence of impurities or contaminants in the GelMA matrix, or the specific protocol used to prepare the membrane can affect the diffusion rate of these molecules through the GelMA membrane. Other biorelevant molecules can have a different charge, backbone or side chains that could possibly interact with GelMA. To gain a better understanding of the interaction between these molecules and the GelMA hydrogel, additional experiments using different types of hydrogels or modified chemical compositions of the hydrogel can be performed, as well as including different biorelevant permeability markers or molecules that have a different chemical composition and properties (e.g. short-chain fatty acids (SCFAs), various pharmaceutical compounds, etc.). For further studies, we decided to continue with the high DS and vary the concentration between 5, 10 and 15 w/v% because high DS was expected to provide optimal mechanical properties, as further evaluated in section 3.2.

### 3.2. GelMA hydrogels exhibit physiologically relevant mechanical stiffness

Recent investigations have underscored the critical role of GelMA concentration in modulating the physicochemical characteristics of hydrogels, and, in turn, dictating their interactions with diverse cell types^47^. A notable trend discerned from these studies is the direct correlation between increasing GelMA concentrations and the enhancement in both the hydrogel’s storage modulus (G’) and adhesive strength. Wu et al. embarked on an in-depth exploration examining the ramifications of GelMA substrate mechanical properties on the outgrowth exhibited by PC12 neuronal cell lines. Their research spanned an array of GelMA-derived hydrogels, with concentrations ranging from 5-30% (stiffness or young’s modulus between 3-180 kPa), hereby providing insights into the dependency of cellular responses on substrate stiffness ^23^. For this reason, we researched the mechanical characteristics of different concentrations of GelMA with a high DS.

Our results in Table 2 show that for all tested GelMA concentrations, the storage modulus is at least 3 times as high as the loss modulus (G”), suggesting that the material is elastic. A higher w/v% of the hydrogel resulted in a higher stiffness of the material. For G’ and G’’, there were significant differences between 5 and 15 w/v% (** p<0.01). The G’ is 1.12 ± 0.17, 6.01 ±1.27, 9.88 ± 1.55 kPa for 5, 10 and 15 w/v% respectively, indicating the disks are soft matrices and rheological values are within normal values for the intestinal lamina propria, whose storage modulus is around 0.5 and 1 kPa^48–50^. According to Deptuła et al., *ex vivo* human intestinal samples are characterized by a stiffness of G’ = 1.52 kPa in the case of healthy tissue and G’ = 9.67 kPa in the case of cancerous tissue ^51^. The measured values are in line with the observation of Gjorevski et al., which suggests 1.3 kPa as the ideal scaffold stiffness for intestinal stem cells (ISC) and organoid cultures ^52^. Soft tissue in general exhibits a storage modulus (G’) from 0.1 kPa to 1 MPa, and all the concentrations result in a hydrogel that is in that range, thus is also suitable as scaffold material for other tissues ^53^.

**Table 2.** Storage modulus (G’), loss modulus (G”), Gel fraction (%) and Mass swelling ratio, measured on 5,10 and 15 w/v% GelMA hydrogel disks (n=4 except for gel fraction and mass swelling ratio of 15w/v% where n=3, values depicted are the average ± standard deviation, data for gel fraction and mass swelling ratio of the 5 w/v% hydrogel disks is not available).

| | $G'$ (kPa) | $G''$ (kPa) | Gel fraction (%) | Mass swelling ratio |
| --- | --- | --- | --- | --- |
| 5 w/v% | $1.12 \pm 0.17$ | $1.44 \pm 0.02$ | NA | NA |
| 10 w/v% | $6.01 \pm 1.27$ | $1.80 \pm 0.28$ | $79.06 \pm 7.95$ | $15.94 \pm 1.81$ |
| 15 w/v% | $9.88 \pm 1.56$ | $2.38 \pm 0.38$ | $89.38 \pm 2.25$ | $7.64 \pm 0.21$ |

Going more in depth on the storage modulus results, previous reports have shown the significance of GelMA concentration in influencing the physicochemical properties of resulting hydrogel and thereby its interaction on different cell types. Studies revealed that when the GelMA concentration was increased, the storage modulus of GelMA hydrogel also increased gradually along with its adhesive strength (reviewed by Kurian and colleagues ^47^). Here, we found that the G’ of the 10 w/v% is 5.4 times higher compared to the 5 w/v% condition, and the 15 w/v% is 8.8 times the 5w/v%, hereby confirming the trend found by previous reports.

The mass swelling ratio is an indicator of how tightly polymer networks are polymerized, with more considerable swelling showing a greater freedom between polymer chains. The results (Table 2) indicate that 10 w/v% hydrogels experience non-significantly higher swelling capacity compared to 15 w/v% hydrogels (p=0.057, M-W U). As for gel fraction, no significant difference between 10 and 15w/v% was observed (p=0.052, M-W U). The gel fraction values reflect which samples were most consistently crosslinked, indicating here that 15 w/v% exhibited a higher degree of polymerization compared to the 10 w/v% hydrogels. No data is presented on the gel fraction and mass swelling ratio of the 5 w/v% hydrogels because of being too brittle and breakage after freeze-drying.

From these results, we can conclude that the storage modulus from all three concentrations is in a representative range to mimic the intestinal tissue, and soft tissue in general. As a next step, we seeded intestinal cells on the scaffold.

### 3.3. GelMA scaffold enables long-term cell adherence, viability and 3D-dome formation

To mimic enterocytes, the human carcinoma-derived immortalized Caco-2 cell line is widely used in *in vitro* models to predict absorption, study permeability and diffusion of compounds through the epithelium and describe mechanisms of host-microbe interactions. After seeding, Caco-2 cells spontaneously differentiate to form a confluent monolayer of polarized cells, structurally and functionally resembling the small intestinal epithelium and expressing several morphological and functional properties characteristic of small bowel enterocytes^54–58^

A preliminary test, where Caco-2 cells were seeded on the hydrogel disks for 72h, showed the cells adhered to the GelMA based on resazurin assay results, as well as corresponding brightfield images (Fig. S6). The lower fluorescence of cells on the 10 w/v% GelMA was likely due to damaging the cells on the edge of the disk when transferring the disk with forceps to a new plate to perform the fluorescence assay. Based on this, the experimental set-up was adjusted by placing the disks on custom 3D-printed hydrogel inserts. The results of the longer-term experiments, displayed in Figure 3, show that the metabolic activity of Caco-2 cells on the hydrogel disks was significantly different between 5-10 w/v% on day 4 after seeding (* p=0.033), suggesting that the cells adhered slightly less on the 5 w/v%. On day 7 after seeding, significant differences in metabolic activity were found between 10 and 15 w/v% and the control and 15 w/v% (* p = 0.033 and * p=0.038 respectively). Interestingly, on day 13, the cell metabolic activity seems to have reached a plateau, on the GelMA as well as on the control. On day 13 and 16, no significant differences were found. On day 20, there was a significant difference in metabolic activity between the control and 5 w/v% hydrogel (** p=0.0063). Fluorescence values increased from day 23 and reached a new plateau, with values indicating the viability of the cells remained stable until the end day 34 (Fig. S5). The viability graph shows the same trend for cells grown on hydrogel disks and the control. On the confocal images of Caco-2 cells on the hydrogel disks (taken after 30 days of culture), it can be observed that the cells form a uniform monolayer on the control as well as the hydrogel membranes, with no 3D dome-formation. Altogether, these from the preliminary, as well as the main experiment results show that Caco-2 cells can adhere to GelMA hydrogel and able to form a stable culture for up to 34 days.

**Figure 3.**
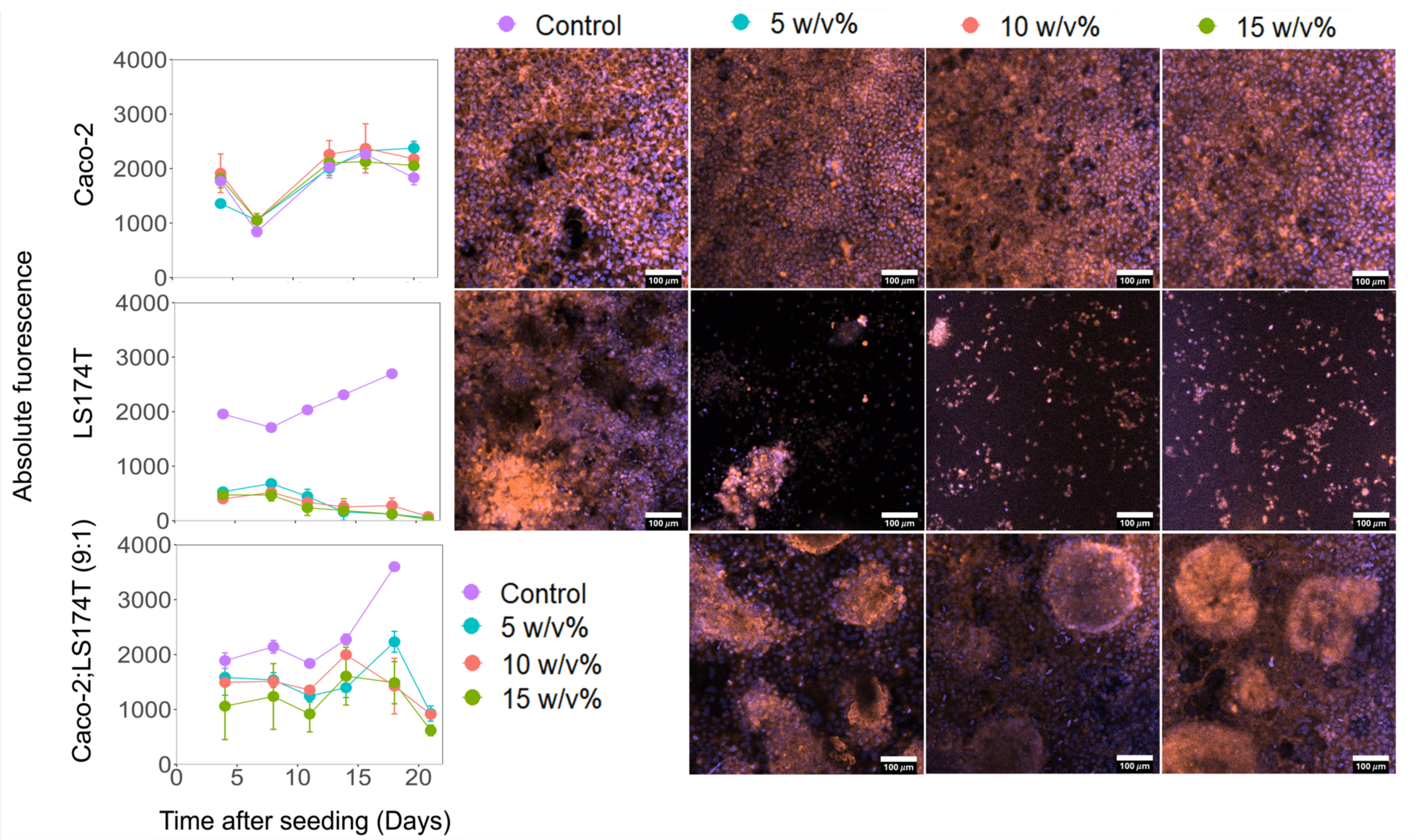
Metabolic activity of the seeded on the intestinal cell lines cells (Caco-2 and LS174T monocultures, and co-cultured) over time on 5 (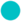), 10 (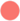) and 15 (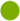)w/v% GelMA disks and control (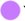) (depicted with dot plots (average ± standard deviation), x-axis shows days after seeding, y-axis shows the absolute measured fluorescence of resorufin, metabolic activity was normalized according to surface area). The metabolic activity is shown until day 20 for Caco-2 cells, and until day 21 for the LS174T monoculture and Caco-2:LS174T co-culture. The metabolic activity of Caco-2 cells on the disks was measured after the dates depicted here and can be found in Fig S5. Next to the metabolic activity plots, confocal images of the cellular morphology of intestinal cells on GelMA disks at the end of the experiment for the mono- and co-cultures, per w/v % of GelMA (Red = Phalloidin-rhodamine staining for exoskeleton; Blue = DAPI staining for cell nuclei). For co-culture and the LS174T monocultures, the experiments were stopped after 24 days since cell seeding. For the Caco-2 cells, the experiment was stopped 34 days after seeding. The confocal images represent the state of the cell right at the point where the experiment was stopped and the cells were fixated, permeabilized and stained.

On the metabolic activity of LS174T cells on the control and hydrogel disks, no significant differences were found in between the concentrations, however, for all days after seeding, there were significant differences between all three membranes and the control (***, p<0.001, ANOVA (n=3)). The confocal images after 24 days of culture show that the LS174T cells did not form a monolayer on the hydrogels and formed small groups of cells rather than a confluent monolayer, which was observed in the control condition. This is confirmed when taking into account brightfield images of the LS174T cells on the disks taken 13 day after seeding, where domes and the presence of single cells can be observed, with no visible cells in between the dome structures. Furthermore, whereas domes can be observed on the hydrogels after 13 days, these domes have mostly disappeared by day 24 where we observe a more sparse distribution of cells on the GelMA (Fig. S3). Furthermore, whereas domes can be observed on the hydrogels after 13 days, these domes have mostly disappeared by day 24. We hypothesize that the cell culture treatment involves exposing a polystyrene microplate to a plasma gas to modify the hydrophobic plastic surface to a hydrophilic surface, where cells can attach to and multiply. This surface does not reflect the *in vivo* situation, where cells adhere to an ECM-like matrix and where the mucus-producing cells do not form a uniform layer, but rather are dispersed in the epithelial cell layer in a ratio of around 10% compared to all other intestinal cell types (i.e., enterocytes, M-cells, enteroendocrine cells, Paneth cells). Thus, our hypothesis is that the physiological GelMA matrix, which exhibits many features like an ECM, provides a more physiological environment for the LS174T cells, and this results in the LS174T cells behaving more like *in vivo* mucus-producing cells.

For the co-culture of the two cell types. Caco-2 and LS174T in a 9:1 ratio differences in metabolic activity between the concentrations were detected on day 8 between control and 15w/v% (*p = 0.033), on day 11 between control and 15 w/v% (*** p = 0.00096) and control and 10 w/v% (* p = 0.041), and on day 14 between control and 5 w/v% (* p = 0.080). At day 18 there were significant differences between all three membranes and the control (control-5 w/v%** p=0.0042; control-10 w/v%***p=0.00019; control-15 w/v% *** p = 0.00024), with the cells seeded on the control having a higher metabolic activity compared to the cells on the hydrogel disks. Interestingly, looking at the confocal images of the co-cultured cells on the hydrogel (Figure 3) (taken after 24 days of co-culture), there is a formation of randomly distributed dome-like structures when combining the two cell types. The domes are typical of transporting epithelial cultures and can typically be seen in differentiated Caco-2 cells and cells present in the domes possess a high degree of maturation ^59^. As it has previously also been hypothesized that the domes are primarily formed by mucus-producing cells, although in this case this cannot be confirmed by the confocal images since there was no staining for MUC2. Here, we also observed dome formation in the control, however, the cells detached before the experiment was stopped, 17 days after seeding. Brightfield images (Fig S4) taken after 13 days of culture, confirm the presence of dome-like structures on both the control and the hydrogel disks. For future research, we suggest repeating this experiment and perform immunostaining for additional markers such as mucus (MUC2), villin (VIL1) and zona occludens-1 (ZO-1) to also provide insight into the status of monolayer differentiation as well as confirm the presence of LS174T cells on the hydrogel in the co-culture at the end. The co-culture of enterocytes and mucus-producing cells has been well established and characterized on Transwell membranes, showing the presence of mucus layer with a physiologically relevant thickness, produced by the LS174T cells^16,60^. The focus of this paper is mainly on basic cell culture parameters (i.e., viability, metabolic activity) and material characterization, and it is clearly shown that Caco-2 cells and the co-cultured cells can adhere to GelMA long-term.

To conclude, significant differences for the metabolic activity of cells seeded on the hydrogels between GelMA concentrations were found on several days for the different cell types. However, overall, this data does not provide a straightforward answer to which concentration supports better cell adhesion and proliferation. We can however conclude that the metabolic activity of the Caco-2 cells on the hydrogels is not significantly different to the control. For the LS174T cells, the cells on the control consistently showed a significantly higher metabolic activity compared to the GelMA, with almost no differences in metabolic activity in between the GelMA concentrations. And finally, especially towards day 16 after seeding, the co-culture cells showed significantly higher metabolic activity on the control compared to the cells on the GelMA disks, with no significant differences in between the GelMA concentrations in terms of metabolic activity. It is thus not straightforward concluding which w/v% of GelMA is a more favorable scaffold for intestinal epithelial cells, however, since the 5 w/v% is more fragile in terms of practical handling, the more optimal concentrations are 10 and 15 w/v%.

An important note here is that a key component of any in vitro model is a source of cells. Cancer cell lines are easy to grow, allow – in most cases, but not always – a direct comparison of experimental results, and are widely used to study molecular mechanisms of (tumor) cell biology^61^. However, they differ metabolically from enterocytes and mucus-producing cells found *in vivo*. A future perspective is to grow patient-specific primary or adult stemcell-derived organoid cultures on the GelMA scaffolds, such that one can model interindividual differences in host-microbe interactions and go towards more precision medicine approaches to better understand diseases like colorectal cancer and inflammatory bowel disease^62^.

Moreover, it has been observed in studies that exposing Caco-2 cells to the fluidic environment shortened the time required for complete differentiation. The gut-on-a-chip models are a step forward towards more relevant and representative micro-physiological systems and allows researchers to study how factors like peristalsis affect microbial behavior and host responses. Several groups reported differentiation of Caco-2 cells after on-chip culture for considerably shorter time, for example 3 to 5 days (whereas normally, the process takes 14 to 21 days) ^63–65^. However, microfluid models might not always be as representative due to their small scale. Bioreactors, on the other hand, could simulate the dynamic environment of the human gut in a larger scale, hereby also providing larger sampling volumes for later analysis of metabolites, proteins etc. As a future perspective, a tubular GelMA scaffold could be made by molding or 3D-printing. This would make it possible to include a tubular GelMA scaffold in a bioreactor and go towards more physiologically representative *in vitro* models of the intestinal microenvironment ^66^. Another future step would be to co-culture epithelial cells, grown on the GelMA scaffold, with live microbiota (using e.g. a microfluidics platform, bioreactor or transwell system), which would allow the study of host-microbe interactions ^16,67^. In this paper, we show that GelMA is an excellent candidate as scaffold material, with potential to be used at scaffold material for epithelial cells in the lumen.

## 4. Conclusion

We experimentally demonstrated the potential of gelatin methacrylamide (GelMA) as a scaffold material for small intestinal epithelial cells, focusing on its permeability, mechanical strength, and biocompatibility. Our findings show that both high and low degrees of substitution (DS) in GelMA ensure adequate permeability for biorelevant transport markers, making it a promising material for *in vitro* modeling of the gut. The study revealed that GelMA supports the long-term adherence and viability of enterocyte-like epithelial cells and co-cultures with mucus-producing cells, showing its versatility and applicability in replicating intestinal cell environments. The mechanical properties of GelMA, across various concentrations, were found to be within the physiological range for intestinal tissues, highlighting its suitability as a scaffold. To conclude, this research further asserts its potential in precision medicine and more sophisticated *in vitro* models like bioreactors or gut-on-a-chip systems. In summary, our research positions GelMA as a valuable tool in the realm of host-microbe interaction studies, offering a path toward more accurate and translatable research in this vital area.

## 6 Acknowledgements

This work was supported by Fonds Wetenschappelijk Onderzoek België (FWO), project grant nr. 3G062519. We would like to acknowledge Geert Meesen for technical assistance with the confocal microscope performed at the Ghent Light Microscopy Core. We thank Dr. Anna Szabó for guiding and assisting in the synthesis and characterization of GelMA, as well as providing photoinitiator. We thank Dr. Jasper Delaey for providing and characterizing the Low DS GelMA that was used in the permeability experiments. Lastly, we would like to thank Pieter De Clercq, Dr. Ludovica Marinelli and Dr. Annelore Beterams for their support in the cell culture lab.

## 7. Author contributions

M.C.A., T.V.D.W., T.G. and P.D conceptualized, supervised and acquired funding for the research. I.R., M.C.A., T.G. and T.V.D.W. conceptualized and designed the experiments. I.R., K.V., N.R. and G.S. carried out the experiments. I.R. analyzed the data. All authors contributed to data interpretation. I.R wrote the manuscript. All authors revised and approved the manuscript.

## 8. Data availability statement

Additional data supporting the findings of this study are available from the corresponding author upon request.

## 9. Additional information (competing interest statement)

The authors declare no competing interests.

## 10. Supplementary data

**Fig. S1:**
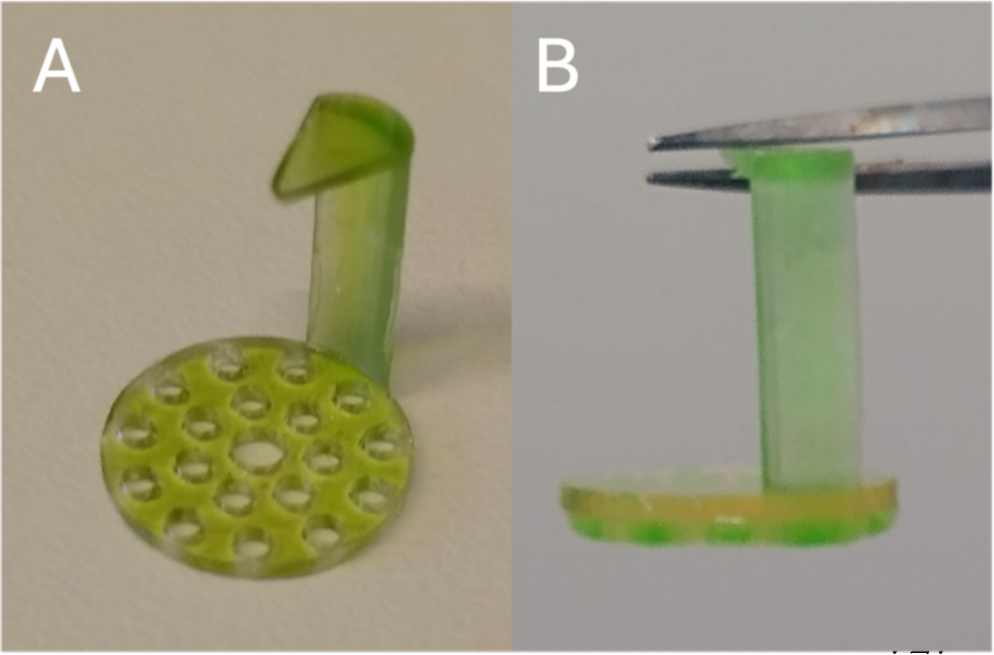
Hydrogel inserts printed in Anycubic translucent green resin with the Anycubic Photon SLA printer. A) Without hydrogel, B) With hydrogel on top of the insert.

**Fig S2:**
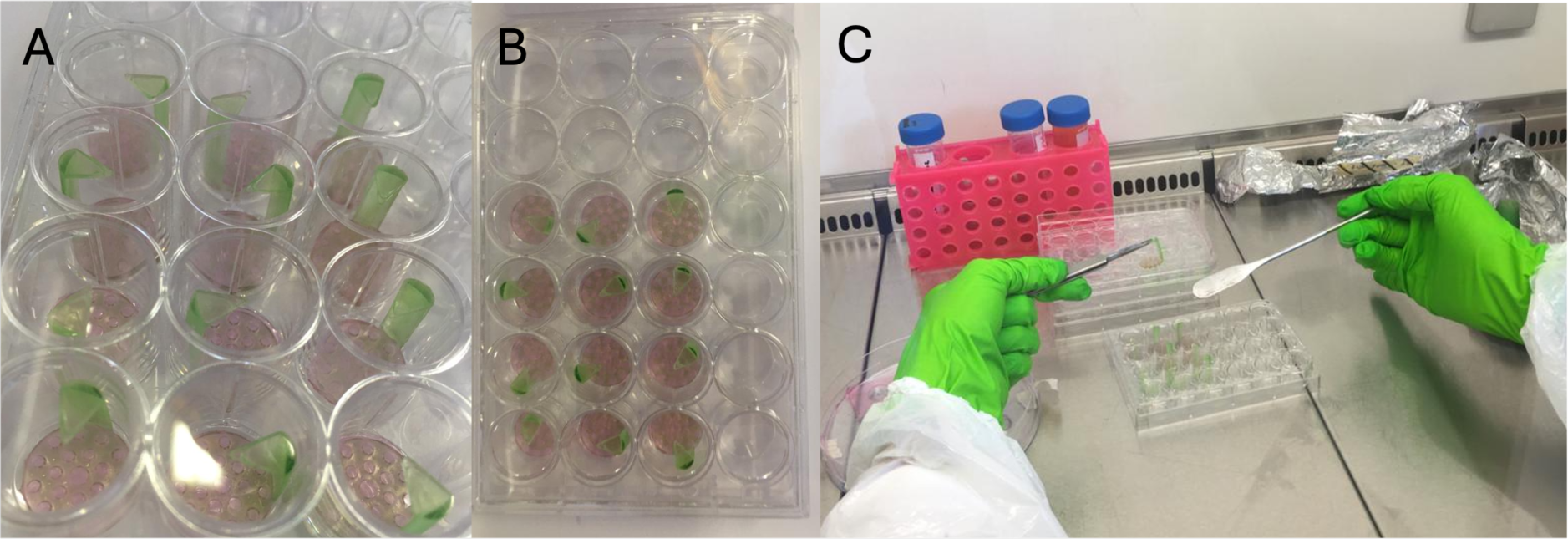
The hydrogel was placed on top of the insert. The insert was designed to have holes on the bottom, providing fresh media to the cells from the bottom but preventing cells from adhering to the plate surface (see Suppl. Fig. 1). After 24h of cell seeding, cell culture media was removed from the wells and the hydrogels were transferred to a new, sterile hydrogel insert. Non-adherent cells were washed away while adhered cells to the GelMA scaffolds were maintained. A and B) inserts with hydrogels in 24-well plate. C) Placing the inserts with hydrogel in the 24-well plate.

**Fig S3:**
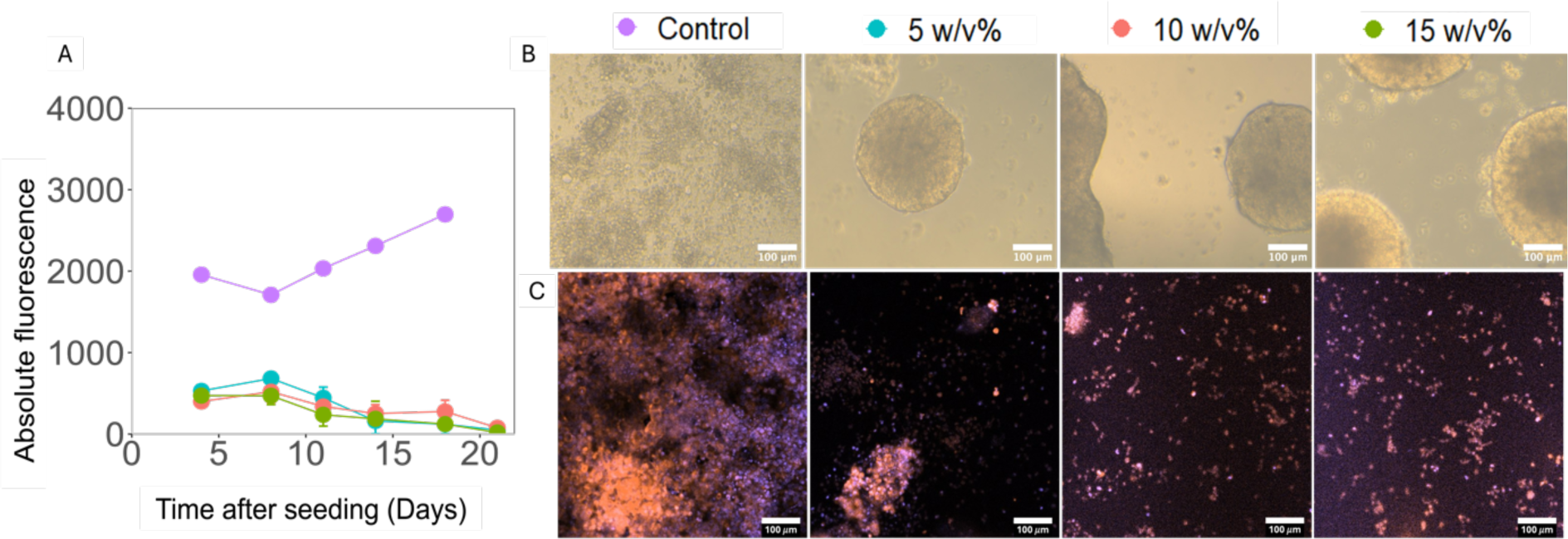
LS174T culture A) Viability over time, measured with resazurin assay. B) Brightfield microscopy images taken 13 days after seeding. C) Confocal images at the end of the experiment (Day 24 after seeding). Red = Phalloidin-rhodamine staining for exoskeleton; Blue = DAPI staining for cell nuclei.

**Fig S4:**
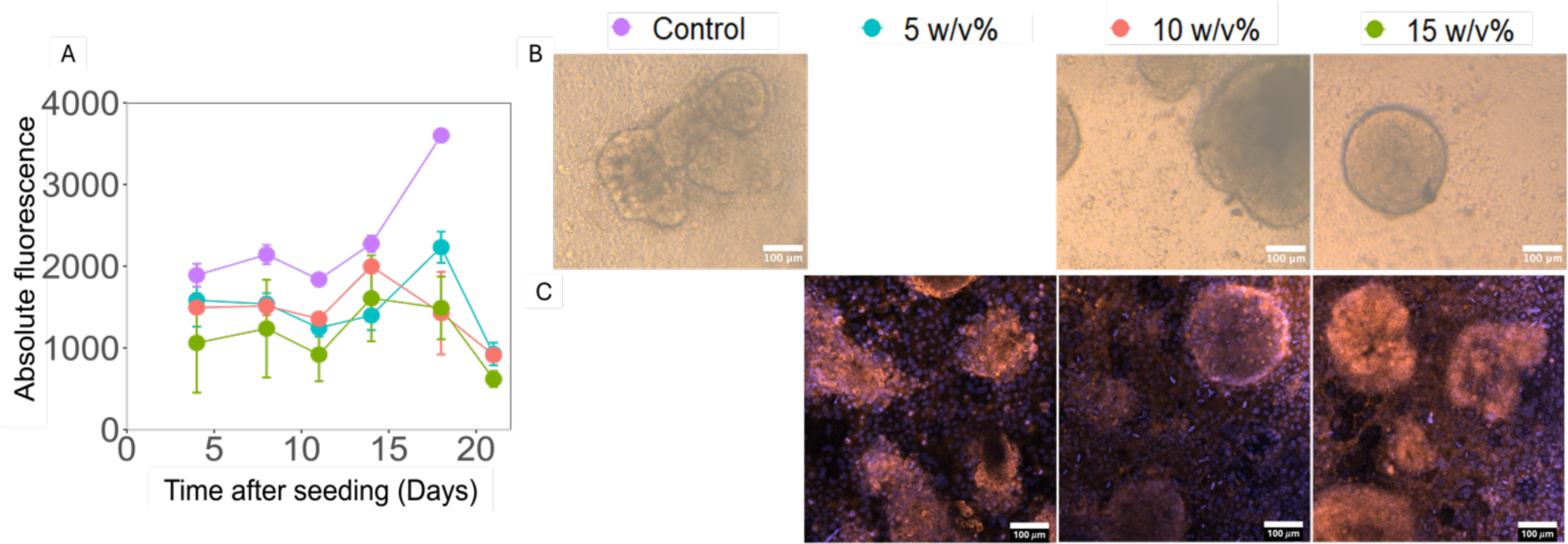
Caco-2 and LS174T co-culture on GelMA. A) Viability over time, measured with resazurin assay. B) Brightfield microscopy images taken 13 days after seeding. No image was taken of the 5 w/v% hydrogel. C) Confocal images at the end of the experiment (Day 24 after seeding). Red = Phalloidin-rhodamine staining for exoskeleton; Blue = DAPI staining for cell nuclei.

**Fig S5:**
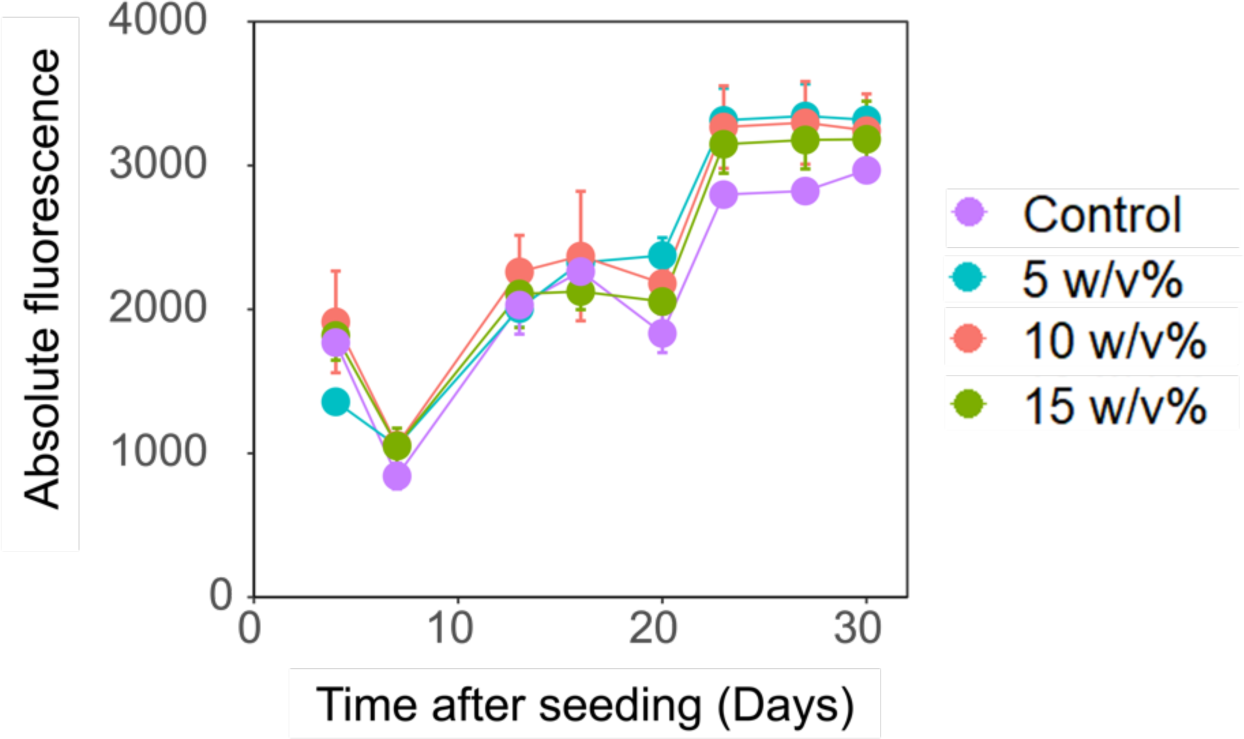
Long-term viability of Caco-2 on GelMA. The experiment with the Caco-2 monoculture was run for 34 days after seeding.

**Fig. S6:**
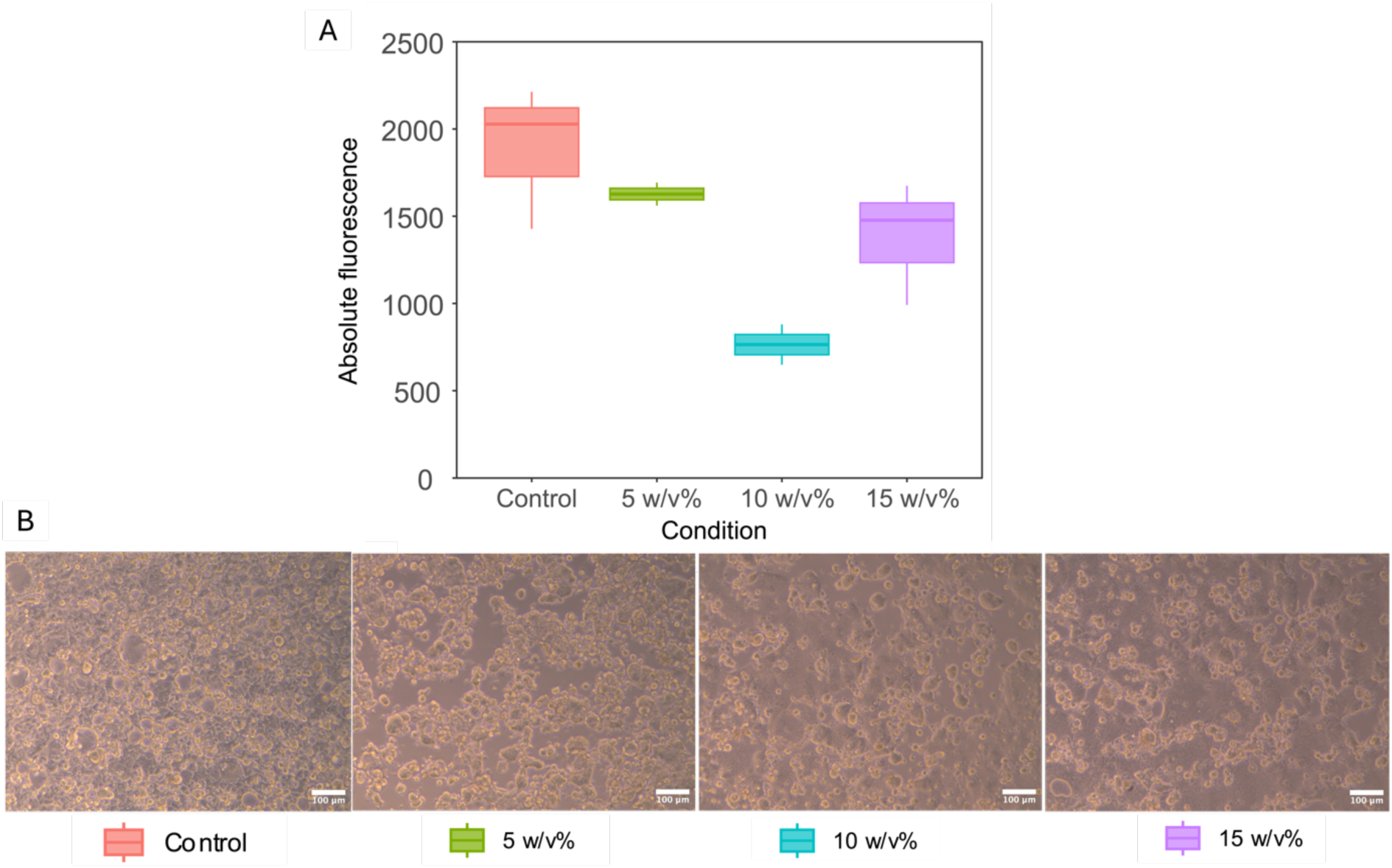
Preliminary results of viability and adherence of Caco-2 culture on GelMA scaffolds (High DS, 5,10 and 15 w/v%, experiment run for 72 hours). A) Resazurin assay data after 72 hours of culture, showing viability of cells on GelMA scaffolds. B) shows corresponding brightfield microscopy images after 72 hours.

## Notes

### Competing Interest Statement

The authors have declared no competing interest.

